# Somatic Mutations Render Human Exome and Pathogen DNA more Similar

**DOI:** 10.1101/322016

**Authors:** Ehsan Ebrahimzadeh, Maggie Engler, David Tse, Razvan Cristescu, Aslan Tchamkerten

**Affiliations:** Department of Electrical Engineering, UCLA, Los Angeles, California, USA; Department of Electrical Engineering, Stanford University, Stanford, California, USA; Department of Discovery Medicine, Merck Research Laboratories, Rahway, New Jersey, USA; Department of Communications and Electronics, Telecom ParisTech, Paris, France

## Abstract

Immunotherapy has recently shown important clinical successes in a substantial number of oncology indications. Additionally, the tumor somatic mutation load has been shown to associate with response to these therapeutic agents, and specific mutational signatures are hypothesized to improve this association, including signatures related to pathogen insults. We sought to study in silico the validity of these observations and addressed three questions. First, we investigated whether somatic mutations typically involved in cancer may increase, in a statistically meaningful manner, the similarity between common pathogens and the human exome. Our study shows that specific common mutagenic processes like those resulting from exposure to ultraviolet light (in melanoma) or smoking (in lung cancer) induce, in the upper range of biologically plausible frequencies, peptides in the cancer exome that are statistically more similar to pathogen peptides than the normal exome. Second, we investigated whether this increased similarity is due to the specificities of the mutagenic process or uniformly random mutations at equal rate would trigger the same effect. For certain pathogens the increased similarity is more pronounced for specific mutagenic processes than for uniformly random mutations and for other pathogens the effects cannot be distinguished. Finally, we investigated whether specific mutational processes result in amino-acid changes with functional relevance that are more likely to be immunogenic. We showed that functional tolerance to mutagenic processes across species generally suggests more resilience to natural processes than to denovo mutagenesis. These results support the idea that recognition of pathogen sequences as well as differential functional tolerance to mutagenic processes may play an important role in the immune recognition process involved in tumor infiltration by lymphocytes.

## Introduction

Recent clinical advances firmly establish the role of immunotherapy (in particular, checkpoint inhibition targetting the CTLA4 and PD1/PD-L1 pathways [1]) in the treatment of cancer. However, the rates of response vary by indication, outlining the important role of identifying the patients most likely to respond [2–5]. In parallel, the analysis of the data in large scale genomic efforts including The Cancer Genome Atlas (TCGA [6]) has identified universal characteristics of the tumor and its environment that ellicit potential recognition by the host immune system. In particular, somatic mutational load as inferred by DNA sequencing [7,8] and cytolytic infiltrate as inferred by RNA sequencing [9] have emerged as hallmarks of an immune-active tumor enviroment. It is thus important to understand the causality and mechanism of action that drives the heterogenous composition of the tumor and its environment and consequently the heterogeneity of response to immunotherapy, in order to select the right patients for treatment, potential combinations, and potential for early intervention.

Multiple recent studies have suggested a strong causal link between the mutational burden of the tumor and clinical response to immunotherapy across multiple indications including Melanoma [10,11], Non Small Cell Lung Cancer [12], Bladder cancer [13] and Colorectal cancer [14]. In these studies, a strong relationship between neoantigen load (the number of mutations with immunogenic potential) and response to immunotherapy has been identified. Importantly, each of these indications are characterized by distinct mutational processes that result in abundant neoantigen load [7, 8]: UV light exposure in Melanoma, smoking in Non Small Cell Lung Cancer, APOBEC activation in Bladder cancer, and MMR defficiency in MSI-h Colorectal cancer. Whether particular mutations or mutational patterns preferentially induce an immunologic phenotype remains an open question [10,11]. However, several hypotheses have recently been put forward, including the presence of mutations in particular genes [15,16], or the presence of a transversion signature related to smoking [12]. In particular, Snyder *et al*. [11] put forward a hypothesis linking cancer exomes with patterns present in common pathogens. Namely, their results with exome analysis of Melanoma patients treated with Ipilimumab, a CTLA4 inhibitor, suggest that somatic mutations in cancer genomes that lead to tetrapeptides similar to those found in common pathogens are more likely to elicit a response to the therapy than common somatic mutations. This association is presumably driven by the innate ability of significant portions of the adaptive immune repertoire to recognize such pathogens.

We took an in-silico approach to evaluate the general validity of this latter hypothesis. Somatic mutations are an inherent natural process related to cell division and aging which in some instances is exacerbated by mutagenic factors. We simulated such mutational processes using mixtures of mutational signatures with empirically derived mixing parameters. We used a simple similarity metric between the mutated exome and common pathogen exomes to estimate changes in overall potential immunogenicity of cancer exomes as compared to the normal exome. We considered simulations of mutational signatures resulting from the mutagenic processes present in most mutated human cancers, namely ultra-violet (UV) light (Melanoma), smoking (Non Small Cell Lung Cancer) and APOBEC activation (Bladder cancer) [7,9]. Our results suggest that, in the upper range of biologically plausible mutation rates, mutational processes enriched in specific alterations resulting from exposure to these common mutagens lead to exome changes that increase the similarity of mutated peptides to peptides of similar sizes originating from common pathogens. These changes are subtle but nevertheless statistically significant and are particularly important in the range of peptide sizes (4–5 amino-acids) that are relevant for epitope presentation in the human MHC presentation mechanism. However, our result also suggest that the increased similarity need not be solely attributed to the specificity of the alteration process. Depending on the pathogen, random mutations (at the same rate) may result in equal increased similarity. These conclusions suggest that mutational processes might act as a mechanism of pressure that models the mutational spectra observed in tumors by increasing recognition from the host immune system.

Opposite to the aforementioned effect that increases the likelihood that a peptide is recognized by the adaptive immune system, an antagonist mechanism of pressure on mutational landscape stems from tolerance by the immune system to natural mutagenic processes. To that extent, we established that exomes across species are generally more resilient, in terms of a functional point of view related to the synonymity of amino-acid changes, to natural mutagenic processes than to denovo mutagenes. In particular, we observe that the functionality of the genetic code (allocation of codons to amino-acids) is more resilient to UV light than smoking mutational processes at a fixed rate. This suggests the possibility that there are different tissue-dependent evolutionary tolerance levels, modulated by the pathogen recognition apparatus in terms of both immune recognition and cancer development, which for example reflect in the much higher mutational loads and immune infiltrate in Melanoma compared to Lung cancer [9].

We thus reveal two key antagonistic mechanisms of pressure that potentially influence the mutational spectra observed in tumors through a differential effect on tolernace. On the one hand similarity to pathogens upper bounds the mutation load by increasing recognition pressure from the host immune system. On the other hand particular mutational process differentially lower bound the mutational load by allowing functional tolerance.

## 1 Methods

We sought to assess whether certain mutational processes result in somatic alterations that increase the similarity of the mutated human exome with selected pathogens. First, we defined a pairwise similarity metric among DNA sequences of varying length. Accordingly, we evaluated the similarity between pathogens and the normal human exome. Second, we simulated mutations resulting from different mutagenic processes acting on the human exome and evaluated the consequent change in similarity of the mutated human exome with respect to the pathogen exomes. Third, we investigated the resiliency of exomes (human exome and model organism exomes) in terms of maintained functionality of the resulting amino-acids and compared the sequences of amino acids of the normal and mutated exomes.

### Data and computing resources

We obtained the human normal exome from GRCh38 http://www.ensembl.org/Homo_sapiens/Info/Index

We considered the following list of model organisms: Mus Musculus (Mouse), Saccharomyces Cerevisiae (Yeast), Felis Catus (Cat), Drosophila Melanogaster (Fruitfly), Caenorhabditis Elegans (Nematode), Xenopus, Danio Rerio (Zebrafish), Cavia Porcellus (Pig), Anolis carolinensis (Anolis). Exomes from these organisms were obtained from http://uswest.ensembl.org/biomart/martview/

We considered the following list of pathogens: Cytomegalovirus (CMV), Dengue virus, Epstein-Barr virus (EBV), Human Herpesvirus 6 (HHV), Human Papillomavirus (HPV), Measles virus, Yellow Fever virus. DNA sequences from these pathogens were obtained from http://www.ncbi.nlm.nih.gov/

We considered simulations of mutational signatures resulting from ultra-violet (UV) light (specific to Melanoma), smoking (specific to Non Small Cell Lung Cancer (NSCLC)), and APOBEC activation (specific to Bladder cancer). These simulations were based on the data from [8, Supplementary information, Table S2] restricted to the set of patients with Melanoma cancer, NSCLC, and Bladder cancer.

For simulations we used Python 2.7.6 (libraries random, numpy, and scipy.stats) and ran programs on a shared server with 8 CPUs and 128GB memory.

## 2 Results

### 2.1 Pathogen DNA vs. human exome and MHC mechanism

To quantify the similarity between a pathogen’s DNA, denoted by *x*, and the human exome, denoted by *y*, we considered the following similarity score. For a given integer *ℓ* ≥ 1, the similarity score, denoted by *sℓ(x, y)*, corresponds to the relative proportion of length-*ℓ* strings in the pathogen DNA that also appear in the human exome at least once, that is 

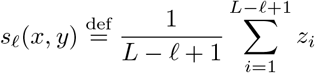

where

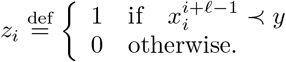

Here L denotes the length of the pathogen’s DNA, 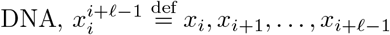 denotes the pathogen’s DNA substring starting at position i and ending at position i + *ℓ —* 1, and “≺” denotes string inclusion.

In particular, *sℓ(x, y)* = 1 corresponds to the case where all length-*ℓ* strings in the pathogen DNA also appear in the human exome and *sℓ(x, y)* &#x003D; 0 corresponds to the case where the pathogen’s DNA and the human exome have no length-ℓ string in common. Observe that *sℓ(x,y)* can be interpreted as the probability that a randomly and uniformly picked length-ℓ string in the pathogen DNA also appears in the human exome.^1^ Because of this we often refer to *sℓ(x,y)* as the matching probability.

In Fig. 1, each curve represents the matching probability *sℓ(x,y)* for a specific pathogen DNA *x* and the normal human exome *y*, for *ℓ* ∈ {9,10,…, 15}. To benchmark these scores we also considered the matching probability with respect to a randomly and uniformly generated pathogen sequence. The average matching probability of a randomly generated pathogen sequence is represented by the “Random” curve in Fig. 1 and turns out to be independent of *L*. This curve is indistinguishable from the 95% confidence interval corresponding to a randomly generated pathogen sequence. Supporting material for Fig. 1 is deferred to the Supplementary Section A.1.

**Figure 1.**
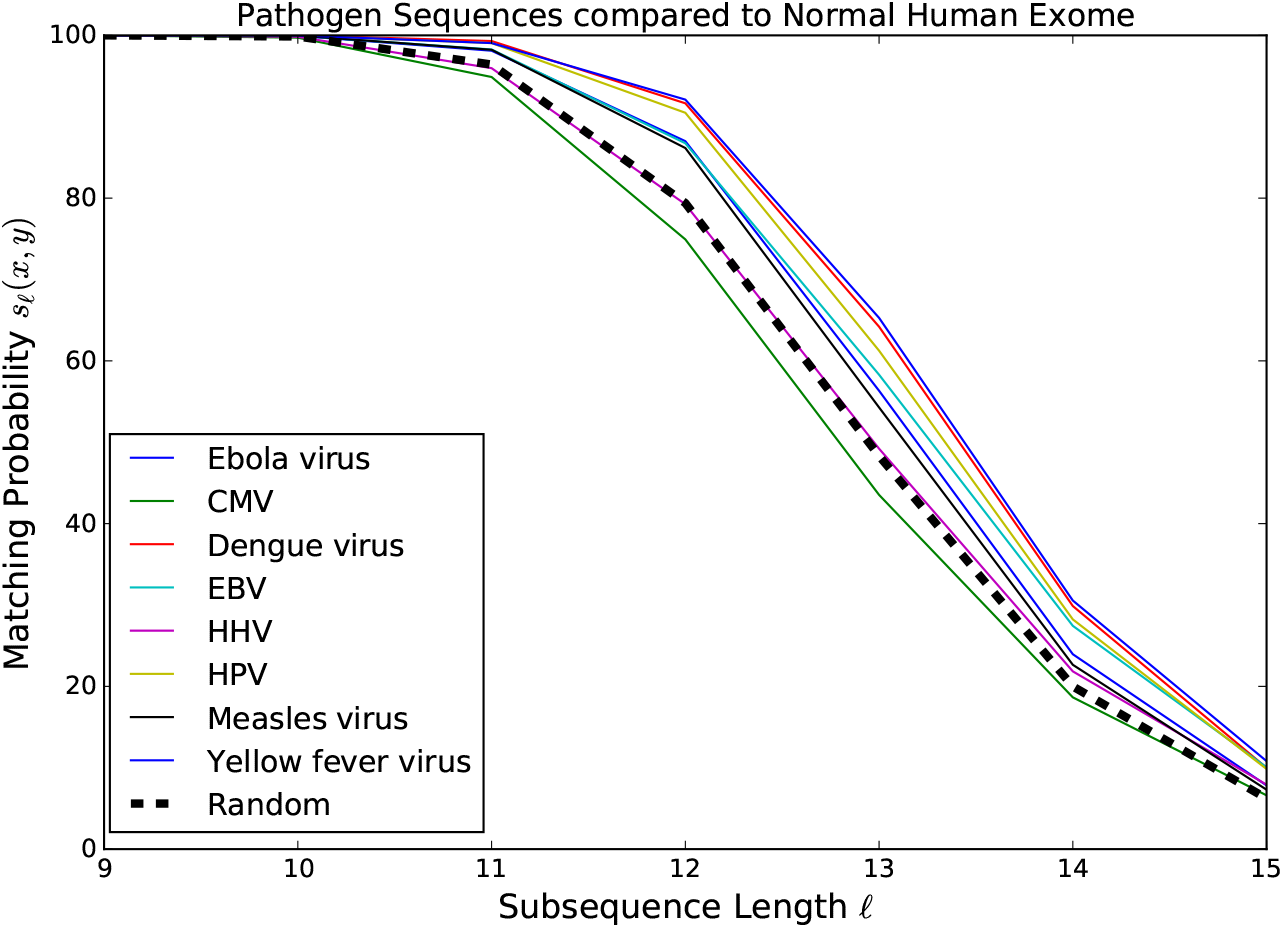
Each curve represents the matching probability (similarity score) *s_ℓ_(x,y)*between a pathogen’s DNA *x* and the human exome *y*, as a function of the subsequence lengths *ℓ*. The “Random” curve refers to the average score of a randomly and uniformly generated “pathogen” DNA sequence.

We observe that the similarity score of a random sequence is generally lower than for pathogen DNA for *ℓ* ≥ 11, except for Dengue virus (*ℓ* ≥ 13) and CMV (always below the random curve). The differences in score across pathogens is maximal at *ℓ*= 12,13 and minimal (zero) at *ℓ* ≤ 10. This latter observation suggests that pathogen and human exome sequences are indistinguishable from each other at a scale corresponding to subsequences of length *ℓ* < 10. Beyond *ℓ*≤10 there is a steep decrease in the similarity score, down to less than 15% for *ℓ*=15. A closer look at the data (see tables in the Supplementary Section A.1) reveals that the sharpest relative drop generally occurs from *ℓ*= 12 to *ℓ*= 13 or from *ℓ*= 14 to *ℓ*= 15. These in-silico observations are in line with the concept that 4–5 amino acids are a sufficiently useful length for the presentation machinery in terms of both diversity of possible sequences (4^12^) and differentiation of self from foreign sequences in the MHC machinery. Namely, this length is strikingly similar to the length of peptides studied in the signature determined by [11].

### 2.2 Impact of mutagens on pathogen DNA and human exome similarity score

To assess whether cancer somatic mutations can make human and pathogen exomes more similar we proceeded as follows. In a first step, we simulated the changes induced to the normal exome by cancer specific mutagens in a probabilistic way. The cancer exomes were generated from the normal exome by using cancer-dependent mixtures of mutational signatures with empirical weights derived from data in [8].^2^ The similarity scores of the normal exome and cancer exome were then computed for each pathogen.

In a second step, we studied the roles of cancer mutation rate and cancer signature. Specifically, we investigated whether random mutations alone, with uniform distribution across mutations, would produce the same results as (typically non-uniform) cancer-dependent mutations, at the same mutation rate.

#### Cancer channel

To formalize our analysis, we used concepts from information theory, in particular related to communications over a noisy channel. To each cancer we associated a transformation, referred to as “cancer channel,” which mimics the effects of the mutagenic processes that are specific to the cancer.^3^ Given a particular cancer c and a mutation rate the cancer channel assigns to each nucleotide *α* the probability ℙ_c_(β|α) of being mutated into nucleotide β. This probability was derived using data from [8, Supplementary information, Table S2] (see Supplementary Section A.2 in this paper).

To obtain a cancer exome ỹ we “passed” the normal human exome y through cancer channel ℙ_c_(·|·) as shown in Fig. 2. Specifically, the cancer exome ỹ was randomly generated from y so that the probability to obtain *ỹ= {ỹ_1_, ỹ_2_,…, ỹ_G_}* from normal exome *y*= {*y*_1_, *y*_2_,…, *y*_G_} was given by

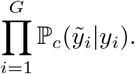

**Figure 2.**
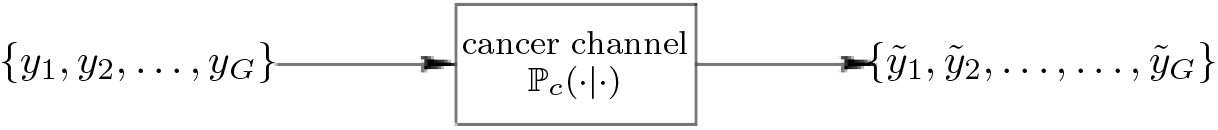
Effects of cancer specific mutations on a normal exome modeled as a cancer channel. Cancer exome ỹ= {ỹ_1_, ỹ_2_,…,ỹ_G_} is obtained from normal exome y= *{y_1_,y_2_,…, y_G_}* through a cancer specific probabilistic transformation ℙ_c_(β|α) which assigns to each nucleotide *α* the probability of being mutated to nucleotide β.

Note that the cancer channel depends on both the mutation rate and the mutation distribution.^4^ The mutation rate was chosen to be equal to 0.001 as it represents a compromise between biological and statistical relevance. It is in the upper range of the mutation rates observed in actual cancer samples [8] and in the lower range for statistical relevance—see next subsection.

#### Normal vs. cancer specific and random mutations

For given pathogen *x* and cancer *c* we performed two tests. In Test 1 we evaluated the statistical significance of the effect of cancer somatic mutations in making human exome more similar to pathogen DNA sequences. In Test 2 we compared cancer somatic mutations and random mutations in making human exome more similar to pathogen DNA sequences.

**Test 1:**For each *ℓ* ∈ {10,11,…, 15} we independently generated 1000 cancer exomes *{ỹ}* from the normal human exome *y* and computed the corresponding similarity scores 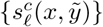. *P*-values were computed for comparing the mean of 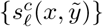 against *s_ℓ_(x,y)* using a one-sided t-test with a null hypothesis that the true mean of 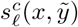 is no larger than *s_ℓ_(x, y)*.

**Test 2:**We replaced the cancer channel by a “random channel” which produced mutations at the same rate (0.001) but in a uniform (1/3,1/3,1/3) manner. For each *ℓ* ∈ {10,11,…, 15} we independently generated 1000 exomes *{ŷ}* by passing the normal human exome *y* through the random channel and computed the corresponding similarity scores {*s_ℓ_(x, ŷ)*}. *P*-values were computed for comparing the mean of 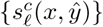 against the mean of 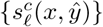 (obtained in Test 1) using a two-sample one-sided t-test with a null hypothesis that the true mean of 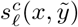 is no larger than the true mean of {*s_ℓ_(x, ŷ)*}.

In Fig. 3, each histogram refers to a particular cancer type. Red bars refer to Test 1 and blue bars refer to Test 2. Bar height represents, for any given *ℓ* ∈ {10,11,…, 15}, the proportion of pathogens for which the *p*-value is ≤ 0.01. Related data can be found in the tables of the Supplementary Sections A.2.1-A.2.8. In these tables, the second
column refers to *s_ℓ_(x, y)*, the third column gives a 95% confidence interval for *s_ℓ_(x, ỹ)*, the fourth column gives the p-value for Test 1 and the fifth column gives the *p*-value for Test 2.

**Figure 3.**
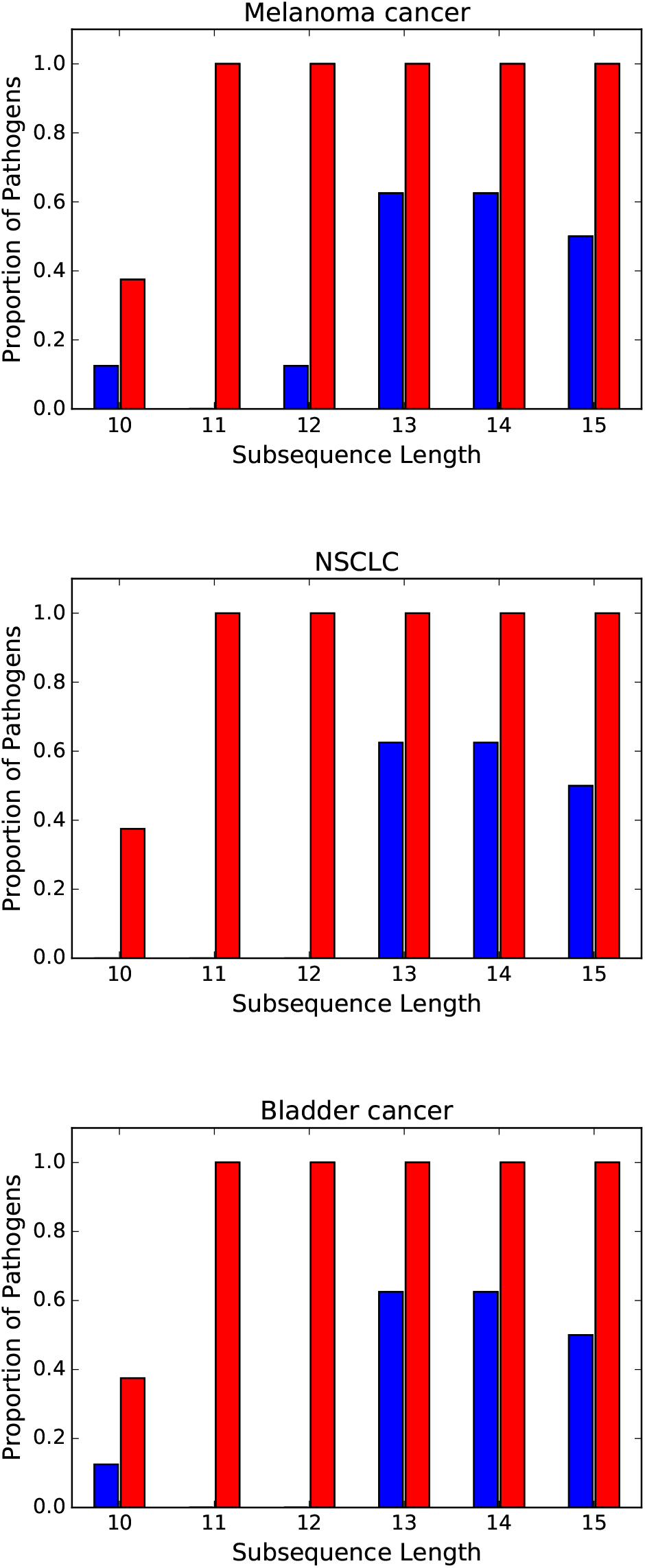
The red bars represent the relative proportions of pathogens whose DNA is more similar to cancer exomes than to normal exome (one-sample t-test results with *p*-value ≤ 0.01). The blue bars represent the relative proportion of pathogens that are more similar to cancer exomes than to exomes with equal mutation rate but uniformly distributed mutations (two-sample one-sided t-test *p*-value ≥ 0.01).

Referring to Test 1 (red bars), we observe that all three mutational processes render the human exome more similar to all pathogen DNA sequences, at all length *ℓ* ∈ {11,12,…, 15}. However, the change is small as it varies by up to 0.5% (see Tables A.2.1-A.2.8, columns 2, 3). Interestingly, whether this change of similarity is due to the specificity of the mutagenic process or random mutations trigger the same effect depends on both the pathogen and the length. By observing the blue bars we see, for instance, that at *ℓ* ∈ {13,14,15}, the change in similarity due to cancer specific mutations is more significant than the change in similarity due to uniformly random mutations for about half of the pathogens. By contrast, at length < 13 the impact of cancer specific mutations cannot be distinguished from uniformly random mutations for most combinations of pathogen and mutation process. We also note a certain consistency across mutational processes. For instance, at length *ℓ* ∈ {13,14}, either the three mutagenic processes render a particular pathogen (Ebola virus, Dengue virus, HHV, HPV, Measles virus) more similar to the normal human exome than uniformly random mutations, or the opposite is true, namely the effect of all three mutagenic processes cannot be distinguished from uniformly random mutations (CMV, EBV, Yellow Fever virus)—see Tables Tables A.2.1-A.2.8, Columns 4 and 5.

### 2.3 Similar resiliency across exomes

In order to compare the resiliency of the model organism exomes with respect to mutagenic processes, we evaluated the error correction capabilities of the genetic code (the codon allocation to amino-acids) for each combination of model exome and mutagenic process.

Referring to Fig. 4, *y*= {*y*_1_,…, *y_L_*} represents a DNA sequence whose corresponding sequence of amino acids is {*a*_1_,…, *a_L_*/_3_}. This DNA sequence is then passed through a given cancer channel ℙ_*c*_(·|·) and results in a cancer sequence *ỹ*= {*ỹ*_1_,*ỹ*_2_,…, *ỹ_L_*} and a corresponding sequence of cancer amino acids {*ã*_1_, *ã*_2_,…, *ã*_*L*/3_}. From {*a*_1_, *a*_2_,…, *a*_*L*/3_} and {*ã*_1_, *ã*_2_,…, *ã*_*L*/3_} we computed the relative proportion of amino acids that were affected, that is

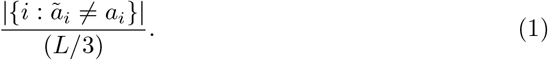

Finally, averaging over all possible realizations of *ỹ* (and therefore over *ã*), we obtained the average error probability

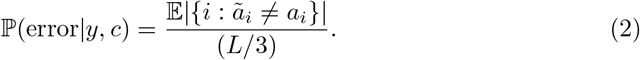

**Figure 4.**
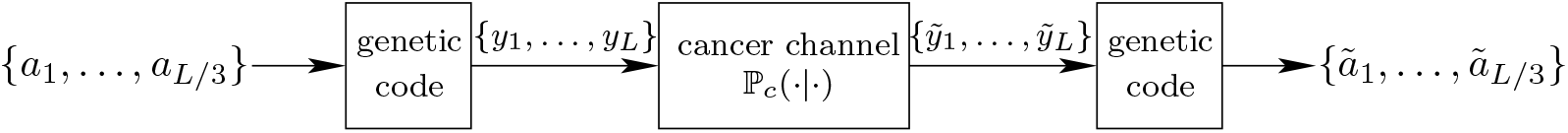
Exome {*y*_1_, *y*_2_, …}, which gives amino acid sequence {*a*_1_, *a*_2_, …}, undergoes specific somatic mutations through transformation ℙ_c_(·|·) and results in {*ỹ*_1_, *ỹ*_2_, …} which, in turn, gives amino acid sequence {*ã*_1_, *ã*_2_, …}.

Fig. 5 represents ℙ(error|*y, c*) for each combination of model organism and cancer at mutation rate 0.001. Details on the computation of ℙ(error|*y, c*) are deferred to the Supplementary Section A.3.

**Figure 5.**
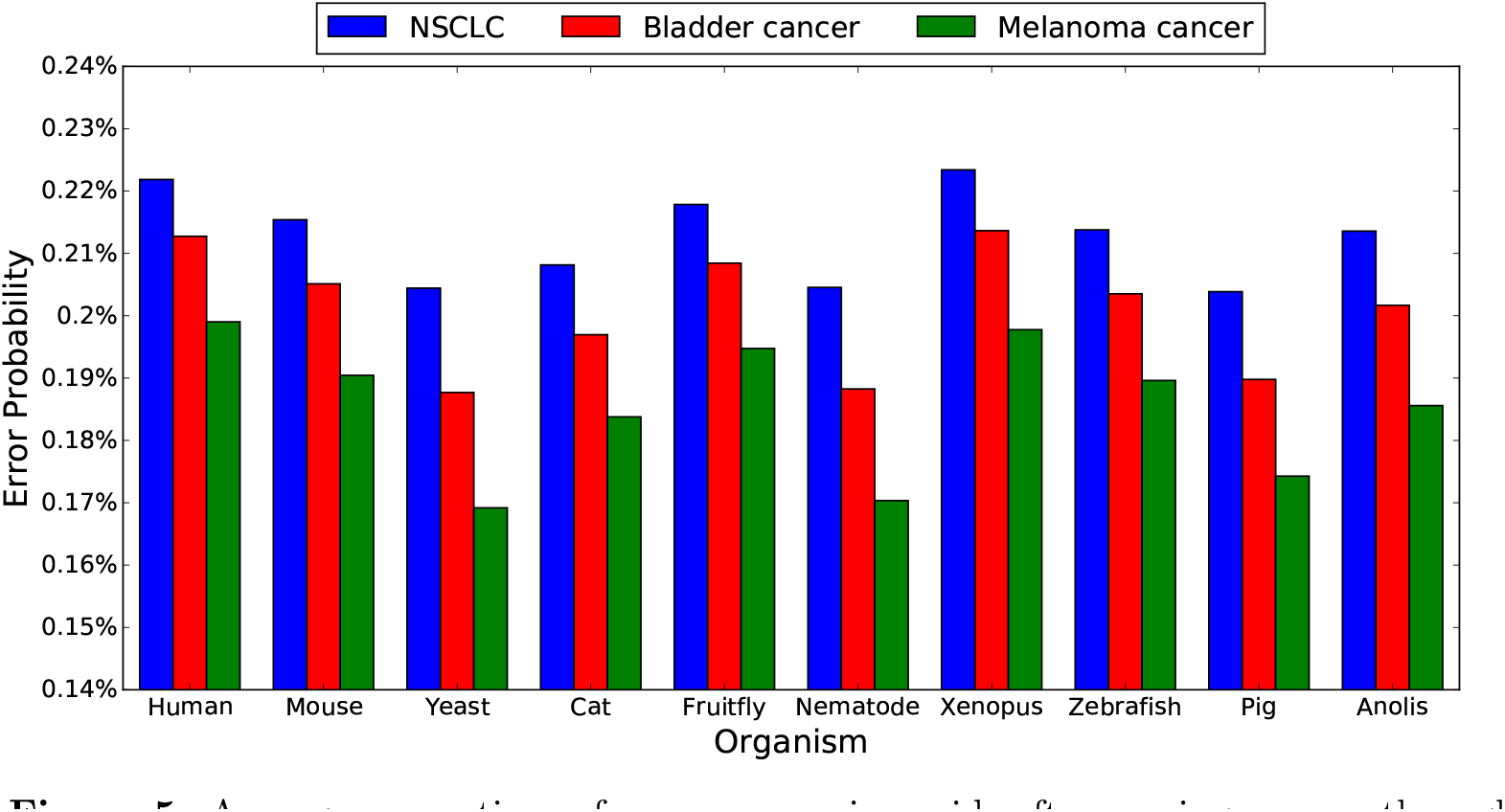
Average proportions of erroneous amino acids after passing exomes through different cancer channels at mutation rate *ρ* = 0:001.

Referring to Fig. 5 we observe that although the proportion of non-synonymous mutations varies across exomes for the three types of mutational processes, it is always lowest for melanoma and maximal for lung. Moreover, this ordering holds irrespectively of the intensity of the mutation rate (see Fig. 6 in the Supplementary Section A.3 which holds for a tenfold mutation rate equal to 0.01). It should be noted that we evaluated the proportions of non-synonymous mutations for several other organisms as well (including the set of pathogens considered in this paper), and could not find one for which the ordering melanoma/bladder/lung did not hold.

**Figure 6.**
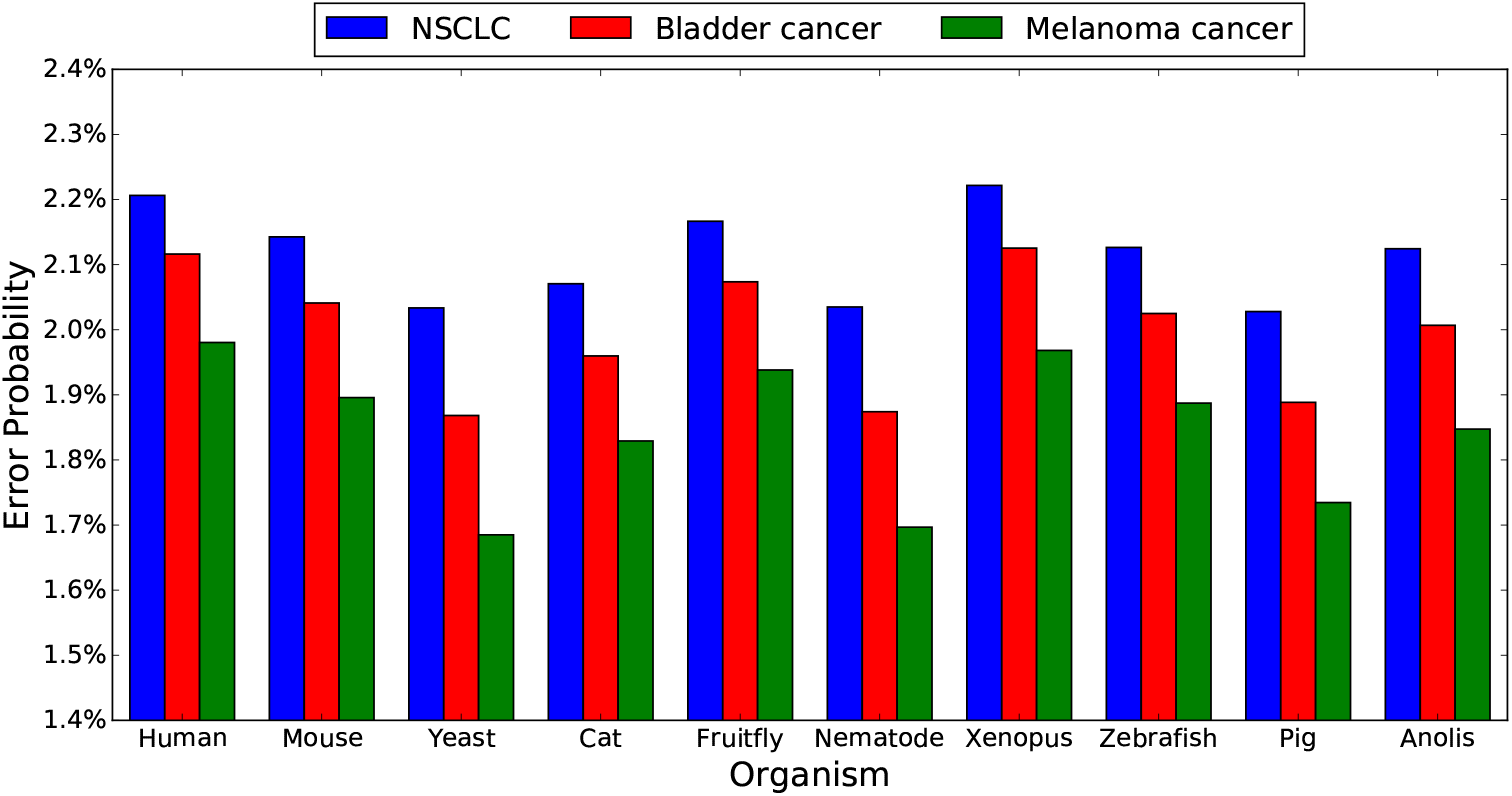
Average proportions of erroneous amino acids after passing exomes through different cancer channels at mutation rate ρ = 0:01.

## 3 Discussion

Our in-silico results show that, in general, the typical stochastic mutational processes encountered in the major cancer indications with abundant neoantigens do appear to shift the peptide distribution of the modified exome in a mutagen-specific manner but universally towards a landscape that appears more similar to pathogenic insult. However, for certain pathogens and length *ℓ* this shift cannot be distinguished from a shift originating from uniformly random mutations, and in this case the mutation rate as opposed to mutation signature is the dominant effect.

We employed large scale simulations to model the random (across space) effect of stochastic mutational processes on the human normal genome. We believe this is a valid approach since the cancer exome available data does suggest that the mutational processes in cancers with large number of mutations affect equally all regions of the exome. We observed that all three major mutational processes considered induce subtle but robust shifts in the measure by which we characterized the similarity between the normal human exome and pathogen sequences, at mutation rates in the upper range of the mutation rates observed in actual cancer samples (0.001). Moreover, the range of peptide lengths where this shift happens aligns with the typical length of peptides presented by the human class I presentation system, suggesting an increased potential for recognition of these types of somatic mutations by a pathogen-trained host immune system.

We also observed that the effect of the considered mutagenic processes on the likelihood of observing a non-synonymous alteration is strikingly different across processes but consistent across the species studied in our framework (human and model organisms). Melanoma/UV light alterations are the least likely to result in aminoacid functional changes, followed by APOBEC-driven alterations and then by smoking alterations, suggesting different error-correcting capabilities of the living exomes towards this various mutagenic insults. This is an attractive observation from an evolutionary perspective: due to universal exposure to sunlight, organisms likely developed similarly universal intrinsic protection from UV light type of modifications to their exomes via the redundancies in the aminoacid codon allocation. Similarly, APOBEC-activation appears to be an universal innate protection mechanism that allows the cell to induce damaging mutations to foreign organisms, while the mutations resulting from tobacco smoking are less likely to have presented evolutionary pressure. In summary, our in-silico approach reveals two competing mechanisms of tolerance pressure on the major mutational processes present in human cancers that modulate the potential immune recognition of alterations at the exome level through pathogen similarity and through functional redundancy; the balance between these mechanisms may significantly contribute to the eventual mutational landscape of advanced cancers.

## 4 Acknowledgements

The authors would like to thank Ka Kit Lam for early discussions on this project.

## A Supplementary information

### A.1 Data for Fig. 1

In the tables below we listed the similarity scores *s_ℓ_(x, y*) of each pathogen *x* against the human exome *y*, as a function of the subsequence length *ℓ*.

| $\ell$ | Ebola virus | CMV | Dengue virus | EBV | HHV |
| --- | --- | --- | --- | --- | --- |
| 9 | 100.0 | 100.0 | 100.0 | 100.0 | 100.0 |
| 10 | 99.94 | 99.71 | 100.0 | 99.94 | 99.83 |
| 11 | 98.11 | 94.88 | 99.30 | 98.26 | 95.96 |
| 12 | 86.99 | 74.92 | 91.65 | 86.74 | 79.24 |
| 13 | 56.32 | 43.52 | 64.19 | 58.29 | 49.20 |
| 14 | 23.94 | 18.64 | 29.87 | 27.43 | 21.84 |
| 15 | 7.82 | 6.60 | 9.98 | 10.05 | 7.91 |

| $\ell$ | HPV | Measles virus | Yellow fever virus | Random |
| --- | --- | --- | --- | --- |
| 9 | 100.0 | 100.0 | 100.0 | 100±0.002 |
| 10 | 99.97 | 99.97 | 100.0 | 100±0.002 |
| 11 | 99.05 | 98.23 | 99.05 | 96.4±0.002 |
| 12 | 90.48 | 86.15 | 92.10 | 79.4±0.002 |
| 13 | 61.24 | 54.27 | 65.28 | 48.3±0.002 |
| 14 | 28.22 | 22.64 | 30.54 | 20±0.002 |
| 15 | 9.84 | 7.29 | 10.81 | 6.2±0.002 |

The column “Random” refers to a 95% confidence interval for the similarity score between a randomly generated pathogen sequence *X*, where each nucleotide is independently and uniformly selected with probability 1 /4, and the normal human exome *y*. To compute this confidence interval we proceeded as follows. The similarity score for a random instance X of length L is given by 

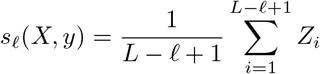

where the *Z_i_*’s are i.i.d. Bernoulli random variables such that

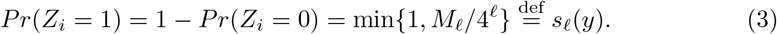

Here *M_ℓ_* denotes the number of distinct length-*ℓ* substrings in the human genome and was computed empirically for *ℓ* ∈ {9,10,…, 15}:

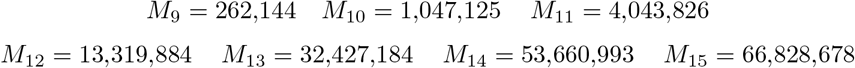

Taking expectation over *X* yields

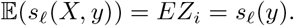

A confidence inteval for *s_ℓ_(X, y)* was computed via Chebyshev’s inequality as follows. We have

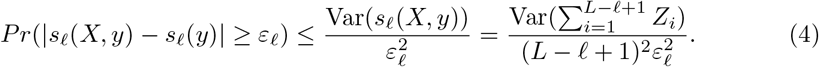

Furthermore, 

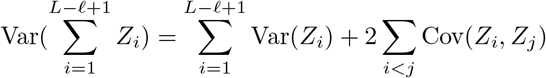

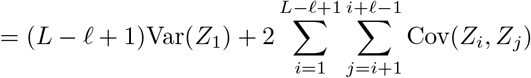

where for the second equality we used the fact that the *Z_i_*’s are identically distributed and that *Z_k_* and *Zj* are independent whenever *j ≥ k* + *ℓ*. Now 

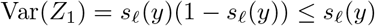

and since the Zi’s are binary random variables 

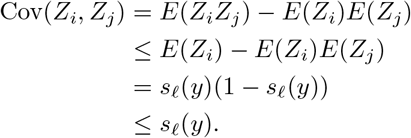

Therefore,

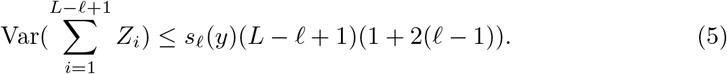

Finally, from (3), (4), and (5) we get 

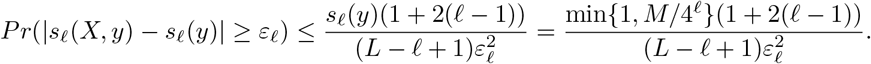

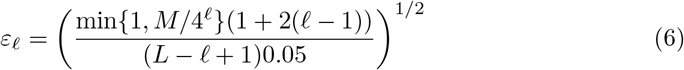

which is below 0.002 for all *ℓ* ∈ {9,10,…, 15} regardless of the pathogen length *L*.

### A.2 Cancer channel and data for Fig. 3

We describe how we obtained cancer channel ℙ*_c_*(·|·) for a given cancer and mutation rate. For each cancer *c* (Melanoma cancer, NSCLC, Bladder cancer) we considered the set *S_c_* of patients in [8, Supplementary information, Table S2] with that cancer. Then, for every mutation *α → β* we empirically computed the average proportion of mutations across patients 

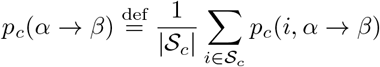

where *p_c_*(*i*,α → β) denotes the proportion of *α → β* mutations among all mutations in patient *i* and was computed from [8, Supplementary information, Table S2]. The probability 

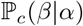

that a nucleotide *α* in the normal exome results in nucleotide *β* in the cancer exome is therefore given by 

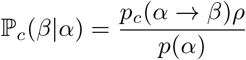

for β ≠ α and 

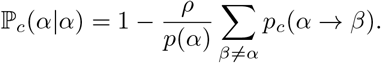

The parameter *ρ* denotes the overall mutation rate and *p(α)* denotes the relative number of nucleotide *α* in the exome and was computed from [8, Supplementary information, Table S2].

**Remark**. *Because in the data from [8, Supplementary information, Table S2] complementary mutations were counted under the same category (e.g., a change from cytosine to tyamine would be treated the same as a change from guanine to adenine), mutation types were considered in pairs. Since the relative proportions of complementary pairs were not given in [8, Supplementary information, Table S2], we made the assumption that they were equal. Hence in the above expression p_c_(i,α → ^* β, i) *actually corresponds to* 

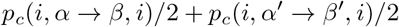

*where* (α’,β’) *is the complementary pair of* (α, β).

The second column in Tables A.2.1-A.2.8 represent *s_ℓ_(x,y)* as a function of *ℓ* (same data as in the tables of the Supplementary Section A.1). The third column represents a 95% confidence interval for 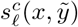 obtained through a standard application of the central limit theorem. This confidence interval is given by 

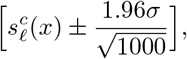

where 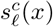 denotes the average of 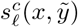 over the 1000 independent trials 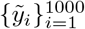 and where *σ* denotes the empirical standard deviation of 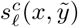. The fourth column gives the *p*-value for Test 1 and the fifth column gives the *p*-value for Test 2.

#### A.2.1 Ebola virus indication

| $\ell$ | Normal | Lung | $p$-value Test 1 | $p$-value Test 2 |
| --- | --- | --- | --- | --- |
| 9 | 100.0 | $99.99 \pm (< 10^{-5})$ | 1.0 | 0.5 |
| 10 | 99.94 | $99.94 \pm 0.0003$ | 1.0 | 0.3328 |
| 11 | 98.11 | $98.16 \pm 0.0015$ | $< 0.01$ | 1.0 |
| 12 | 86.99 | $87.16 \pm 0.0031$ | $< 0.01$ | 0.2037 |
| 13 | 56.32 | $56.58 \pm 0.0045$ | $< 0.01$ | $< 0.01$ |
| 14 | 23.94 | $24.11 \pm 0.0040$ | $< 0.01$ | $< 0.01$ |
| 15 | 7.82 | $7.88 \pm 0.0025$ | $< 0.01$ | $< 0.01$ |

| $\ell$ | Normal | Bladder | $p$-value Test 1 | $p$-value Test 2 |
| --- | --- | --- | --- | --- |
| 9 | 100.0 | $99.99 \pm (< 10^{-5})$ | 1.0 | 0.5 |
| 10 | 99.94 | $99.94 \pm 0.0003$ | 0.9999 | $< 0.01$ |
| 11 | 98.11 | $98.17 \pm 0.0015$ | $< 0.01$ | 0.9999 |
| 12 | 86.99 | $87.16 \pm 0.0031$ | $< 0.01$ | 0.4714 |
| 13 | 56.32 | $56.57 \pm 0.0043$ | $< 0.01$ | $< 0.01$ |
| 14 | 23.94 | $24.10 \pm 0.0038$ | $< 0.01$ | $< 0.01$ |
| 15 | 7.82 | $7.88 \pm 0.0025$ | $< 0.01$ | $< 0.01$ |

| $\ell$ | Normal | Mela | $p$-value Test 1 | $p$-value Test 2 |
| --- | --- | --- | --- | --- |
| 9 | 100.0 | $99.99 \pm (< 10^{-5})$ | 1.0 | 0.5 |
| 10 | 99.94 | $99.94 \pm 0.0002$ | 1.0 | 0.8304 |
| 11 | 98.11 | $98.16 \pm 0.0015$ | $< 0.01$ | 1.0 |
| 12 | 86.99 | $87.15 \pm 0.0032$ | $< 0.01$ | 0.9989 |
| 13 | 56.32 | $56.59 \pm 0.0045$ | $< 0.01$ | $< 0.01$ |
| 14 | 23.94 | $24.12 \pm 0.0039$ | $< 0.01$ | $< 0.01$ |
| 15 | 7.82 | $7.89 \pm 0.0025$ | $< 0.01$ | $< 0.01$ |

#### A.2.2 CMV indication

| $\ell$ | Normal | Lung | $p$-value Test 1 | $p$-value Test 2 |
| --- | --- | --- | --- | --- |
| 9 | 100.0 | $99.99 \pm (< 10^{-5})$ | 1.0 | 0.6082 |
| 10 | 99.71 | $99.73 \pm 0.0002$ | $< 0.01$ | 0.9999 |
| 11 | 94.88 | $95.01 \pm 0.0007$ | $< 0.01$ | 1.0 |
| 12 | 74.92 | $75.18 \pm 0.0011$ | $< 0.01$ | 1.0 |
| 13 | 43.52 | $43.74 \pm 0.0012$ | $< 0.01$ | 1.0 |
| 14 | 18.64 | $18.75 \pm 0.0010$ | $< 0.01$ | 1.0 |
| 15 | 6.60 | $6.63 \pm 0.0006$ | $< 0.01$ | 1.0 |

| $\ell$ | Normal | Bladder | $p$-value Test 1 | $p$-value Test 2 |
| --- | --- | --- | --- | --- |
| 9 | 100.0 | $99.99 \pm (< 10^{-5})$ | 1.0 | 0.5183 |
| 10 | 99.71 | $99.73 \pm 0.0002$ | $< 0.01$ | 0.0255 |
| 11 | 94.88 | $95.02 \pm 0.0007$ | $< 0.01$ | 0.9999 |
| 12 | 74.92 | $75.21 \pm 0.0011$ | $< 0.01$ | 1.0 |
| 13 | 43.52 | $43.76 \pm 0.0012$ | $< 0.01$ | 1.0 |
| 14 | 18.64 | $18.76 \pm 0.0010$ | $< 0.01$ | 1.0 |
| 15 | 6.60 | $6.64 \pm 0.0007$ | $< 0.01$ | 1.0 |

| $\ell$ | Normal | Mela | $p$-value Test 1 | $p$-value Test 2 |
| --- | --- | --- | --- | --- |
| 9 | 100.0 | $99.99 \pm (< 10^{-5})$ | 1.0 | 0.7682 |
| 10 | 99.71 | $99.73 \pm 0.0002$ | $< 0.01$ | 1.0 |
| 11 | 94.88 | $95.00 \pm 0.0007$ | $< 0.01$ | 1.0 |
| 12 | 74.92 | $75.16 \pm 0.0011$ | $< 0.01$ | 1.0 |
| 13 | 43.52 | $43.72 \pm 0.0012$ | $< 0.01$ | 1.0 |
| 14 | 18.64 | $18.73 \pm 0.0010$ | $< 0.01$ | 1.0 |
| 15 | 6.60 | $6.63 \pm 0.0006$ | $< 0.01$ | 1.0 |

#### A.2.3 Dengue virus indication

| $\ell$ | Normal | Lung | $p$-value Test 1 | $p$-value Test 2 |
| --- | --- | --- | --- | --- |
| 9 | 100.0 | $99.99 \pm (< 10^{-5})$ | 1.0 | 0.5 |
| 10 | 100.0 | $99.98 \pm (< 10^{-5})$ | 1.0 | 0.3914 |
| 11 | 99.30 | $99.31 \pm 0.0014$ | $< 0.01$ | 0.9896 |
| 12 | 91.65 | $91.75 \pm 0.0038$ | $< 0.01$ | 0.9918 |
| 13 | 64.19 | $64.40 \pm 0.0060$ | $< 0.01$ | $< 0.01$ |
| 14 | 29.87 | $30.01 \pm 0.0056$ | $< 0.01$ | $< 0.01$ |
| 15 | 9.98 | $10.04 \pm 0.0039$ | $< 0.01$ | $< 0.01$ |

| $\ell$ | Normal | Bladder | $p$-value Test 1 | $p$-value Test 2 |
| --- | --- | --- | --- | --- |
| 9 | 100.0 | $99.99 \pm (< 10^{-5})$ | 1.0 | 0.5 |
| 10 | 100.0 | $99.98 \pm 0.0001$ | 1.0 | 0.8145 |
| 11 | 99.30 | $99.31 \pm 0.0014$ | $< 0.01$ | 0.9694 |
| 12 | 91.65 | $91.75 \pm 0.0036$ | $< 0.01$ | 0.9986 |
| 13 | 64.19 | $64.39 \pm 0.0059$ | $< 0.01$ | $< 0.01$ |
| 14 | 29.87 | $30.01 \pm 0.0057$ | $< 0.01$ | $< 0.01$ |
| 15 | 9.98 | $10.04 \pm 0.0039$ | $< 0.01$ | $< 0.01$ |

| $\ell$ | Normal | Mela | $p$-value Test 1 | $p$-value Test 2 |
| --- | --- | --- | --- | --- |
| 9 | 100.0 | $99.99 \pm (< 10^{-5})$ | 1.0 | 0.5 |
| 10 | 100.0 | $99.98 \pm (< 10^{-5})$ | 1.0 | 0.4457 |
| 11 | 99.30 | $99.30 \pm 0.0013$ | $< 0.01$ | 0.9999 |
| 12 | 91.65 | $91.75 \pm 0.0035$ | $< 0.01$ | 0.9999 |
| 13 | 64.19 | $64.41 \pm 0.0061$ | $< 0.01$ | $< 0.01$ |
| 14 | 29.87 | $30.02 \pm 0.0055$ | $< 0.01$ | $< 0.01$ |
| 15 | 9.98 | $10.05 \pm 0.0037$ | $< 0.01$ | $< 0.01$ |

#### A.2.4 EBV indication

| $\ell$ | Normal | Lung | $p$-value Test 1 | $p$-value Test 2 |
| --- | --- | --- | --- | --- |
| 9 | 100.0 | $99.99 \pm (< 10^{-5})$ | 1.0 | 0.1418 |
| 10 | 99.94 | $99.95 \pm 0.0001$ | $< 0.01$ | 0.7993 |
| 11 | 98.26 | $98.31 \pm 0.0005$ | $< 0.01$ | 1.0 |
| 12 | 86.74 | $86.91 \pm 0.0012$ | $< 0.01$ | 1.0 |
| 13 | 58.29 | $58.49 \pm 0.0016$ | $< 0.01$ | 1.0 |
| 14 | 27.43 | $27.54 \pm 0.0015$ | $< 0.01$ | 1.0 |
| 15 | 10.05 | $10.08 \pm 0.0012$ | $< 0.01$ | 1.0 |

| $\ell$ | Normal | Bladder | $p$-value Test 1 | $p$-value Test 2 |
| --- | --- | --- | --- | --- |
| 9 | 100.0 | $99.99 \pm (< 10^{-5})$ | 1.0 | 0.2185 |
| 10 | 99.94 | $99.95 \pm 0.0001$ | $< 0.01$ | 0.1288 |
| 11 | 98.26 | $98.31 \pm 0.0005$ | $< 0.01$ | 0.9999 |
| 12 | 86.74 | $86.92 \pm 0.0012$ | $< 0.01$ | 1.0 |
| 13 | 58.29 | $58.50 \pm 0.0016$ | $< 0.01$ | 1.0 |
| 14 | 27.43 | $27.55 \pm 0.0014$ | $< 0.01$ | 1.0 |
| 15 | 10.05 | $10.09 \pm 0.0012$ | $< 0.01$ | 1.0 |

| $\ell$ | Normal | Mela | $p$-value Test 1 | $p$-value Test 2 |
| --- | --- | --- | --- | --- |
| 9 | 100.0 | $99.99 \pm (< 10^{-5})$ | 1.0 | 0.0412 |
| 10 | 99.94 | $99.95 \pm 0.0001$ | $< 0.01$ | 0.9878 |
| 11 | 98.26 | $98.31 \pm 0.0005$ | $< 0.01$ | 1.0 |
| 12 | 86.74 | $86.90 \pm 0.0012$ | $< 0.01$ | 1.0 |
| 13 | 58.29 | $58.47 \pm 0.0016$ | $< 0.01$ | 1.0 |
| 14 | 27.43 | $27.52 \pm 0.0014$ | $< 0.01$ | 1.0 |
| 15 | 10.05 | $10.07 \pm 0.0011$ | $< 0.01$ | 1.0 |

#### A.2.5 HHV indication

| $\ell$ | Normal | Lung | $p$-value Test 1 | $p$-value Test 2 |
| --- | --- | --- | --- | --- |
| 9 | 100.0 | $99.99 \pm (< 10^{-5})$ | 1.0 | 0.2959 |
| 10 | 99.83 | $99.84 \pm 0.0002$ | $< 0.01$ | 1.0 |
| 11 | 95.96 | $96.07 \pm 0.0008$ | $< 0.01$ | 1.0 |
| 12 | 79.24 | $79.48 \pm 0.0018$ | $< 0.01$ | 1.0 |
| 13 | 49.20 | $49.45 \pm 0.0021$ | $< 0.01$ | $< 0.01$ |
| 14 | 21.84 | $21.98 \pm 0.0018$ | $< 0.01$ | $< 0.01$ |
| 15 | 7.91 | $7.97 \pm 0.0010$ | $< 0.01$ | $< 0.01$ |

| $\ell$ | Normal | Bladder | $p$-value Test 1 | $p$-value Test 2 |
| --- | --- | --- | --- | --- |
| 9 | 100.0 | $99.99 \pm (< 10^{-5})$ | 1.0 | 0.5 |
| 10 | 99.83 | $99.84 \pm 0.0002$ | $< 0.01$ | 0.9739 |
| 11 | 95.96 | $96.08 \pm 0.0008$ | $< 0.01$ | 1.0 |
| 12 | 79.24 | $79.49 \pm 0.0018$ | $< 0.01$ | 0.9999 |
| 13 | 49.20 | $49.45 \pm 0.0023$ | $< 0.01$ | $< 0.01$ |
| 14 | 21.84 | $21.98 \pm 0.0018$ | $< 0.01$ | $< 0.01$ |
| 15 | 7.91 | $7.97 \pm 0.0010$ | $< 0.01$ | $< 0.01$ |

| $\ell$ | Normal | Mela | $p$-value Test 1 | $p$-value Test 2 |
| --- | --- | --- | --- | --- |
| 9 | 100.0 | $99.99 \pm (< 10^{-5})$ | 1.0 | 0.8155 |
| 10 | 99.83 | $99.84 \pm 0.0002$ | $< 0.01$ | 1.0 |
| 11 | 95.96 | $96.06 \pm 0.0008$ | $< 0.01$ | 1.0 |
| 12 | 79.24 | $79.47 \pm 0.0020$ | $< 0.01$ | 1.0 |
| 13 | 49.20 | $49.45 \pm 0.0022$ | $< 0.01$ | $< 0.01$ |
| 14 | 21.84 | $21.99 \pm 0.0017$ | $< 0.01$ | $< 0.01$ |
| 15 | 7.91 | $7.97 \pm 0.0011$ | $< 0.01$ | $< 0.01$ |

#### A.2.6 HPV indication

| $\ell$ | Normal | Lung | $p$-value Test 1 | $p$-value Test 2 |
| --- | --- | --- | --- | --- |
| 9 | 100.0 | $99.98 \pm (< 10^{-5})$ | 1.0 | 0.5 |
| 10 | 99.97 | $99.96 \pm 0.0004$ | 1.0 | 0.4823 |
| 11 | 99.05 | $99.05 \pm 0.0016$ | $< 0.01$ | 0.9992 |
| 12 | 90.48 | $90.61 \pm 0.0044$ | $< 0.01$ | 0.0447 |
| 13 | 61.24 | $61.48 \pm 0.0068$ | $< 0.01$ | $< 0.01$ |
| 14 | 28.22 | $28.39 \pm 0.0062$ | $< 0.01$ | $< 0.01$ |
| 15 | 9.84 | $9.91 \pm 0.0042$ | $< 0.01$ | $< 0.01$ |

| $\ell$ | Normal | Bladder | $p$-value Test 1 | $p$-value Test 2 |
| --- | --- | --- | --- | --- |
| 9 | 100.0 | $99.98 \pm (< 10^{-5})$ | 1.0 | 0.5 |
| 10 | 99.97 | $99.96 \pm 0.0003$ | 1.0 | 0.2536 |
| 11 | 99.05 | $99.06 \pm 0.0016$ | $< 0.01$ | 0.9969 |
| 12 | 90.48 | $90.61 \pm 0.0044$ | $< 0.01$ | 0.2479 |
| 13 | 61.24 | $61.47 \pm 0.0067$ | $< 0.01$ | $< 0.01$ |
| 14 | 28.22 | $28.38 \pm 0.0061$ | $< 0.01$ | $< 0.01$ |
| 15 | 9.84 | $9.91 \pm 0.0040$ | $< 0.01$ | $< 0.01$ |

| $\ell$ | Normal | Mela | $p$-value Test 1 | $p$-value Test 2 |
| --- | --- | --- | --- | --- |
| 9 | 100.0 | $99.98 \pm (< 10^{-5})$ | 1.0 | 0.5 |
| 10 | 99.97 | $99.96 \pm 0.0003$ | 1.0 | 0.4093 |
| 11 | 99.05 | $99.05 \pm 0.0016$ | $< 0.01$ | 0.9998 |
| 12 | 90.48 | $90.62 \pm 0.0043$ | $< 0.01$ | $< 0.01$ |
| 13 | 61.24 | $61.50 \pm 0.0069$ | $< 0.01$ | $< 0.01$ |
| 14 | 28.22 | $28.40 \pm 0.0059$ | $< 0.01$ | $< 0.01$ |
| 15 | 9.84 | $9.92 \pm 0.0040$ | $< 0.01$ | $< 0.01$ |

#### A.2.7 Measles virus indication

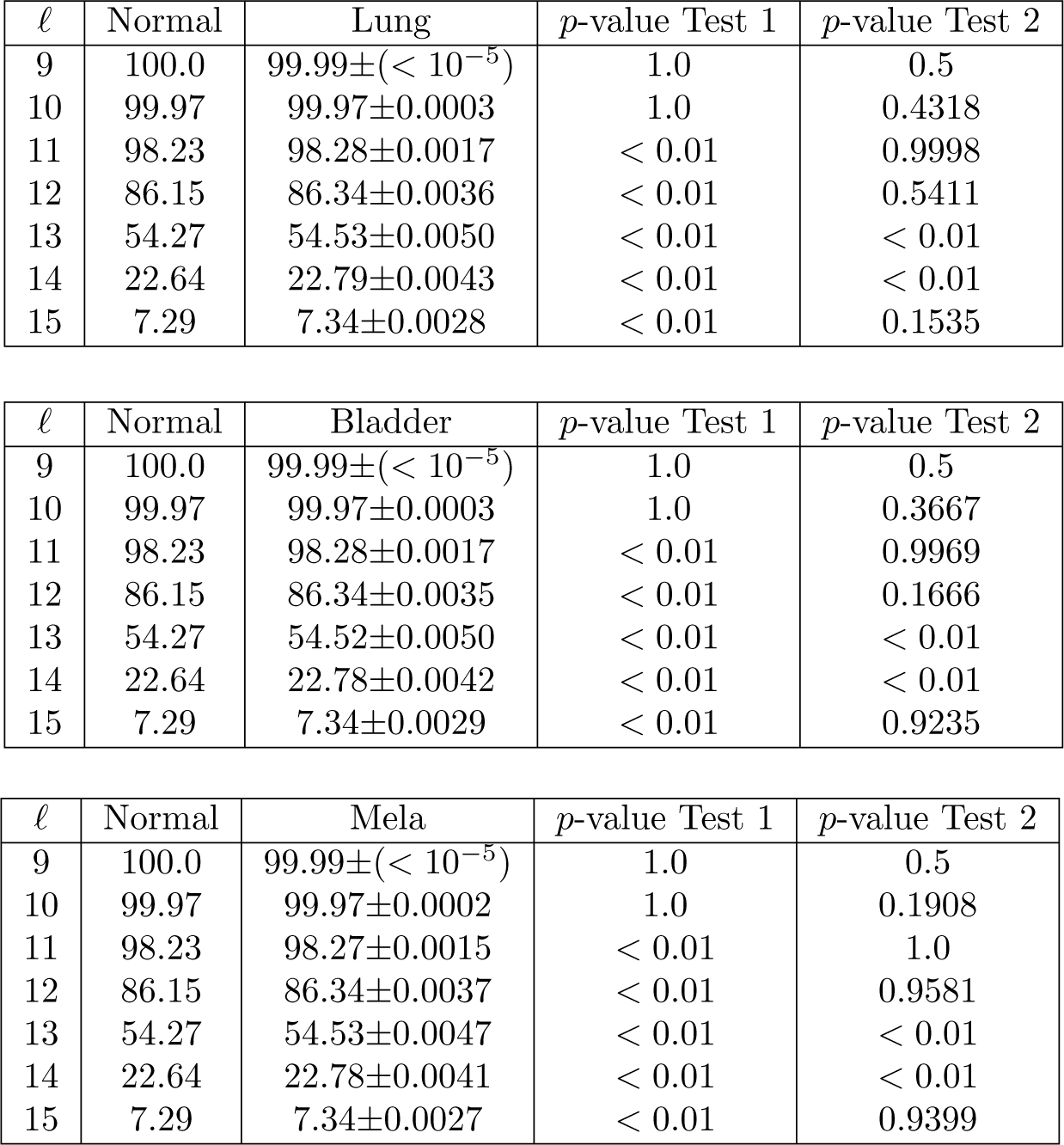

| $\ell$ | Normal | Lung | $p$-value Test 1 | $p$-value Test 2 |
| --- | --- | --- | --- | --- |
| 9 | 100.0 | $99.99 \pm (< 10^{-5})$ | 1.0 | 0.5 |
| 10 | 99.97 | $99.97 \pm 0.0003$ | 1.0 | 0.4318 |
| 11 | 98.23 | $98.28 \pm 0.0017$ | $< 0.01$ | 0.9998 |
| 12 | 86.15 | $86.34 \pm 0.0036$ | $< 0.01$ | 0.5411 |
| 13 | 54.27 | $54.53 \pm 0.0050$ | $< 0.01$ | $< 0.01$ |
| 14 | 22.64 | $22.79 \pm 0.0043$ | $< 0.01$ | $< 0.01$ |
| 15 | 7.29 | $7.34 \pm 0.0028$ | $< 0.01$ | 0.1535 |

| $\ell$ | Normal | Bladder | $p$-value Test 1 | $p$-value Test 2 |
| --- | --- | --- | --- | --- |
| 9 | 100.0 | $99.99 \pm (< 10^{-5})$ | 1.0 | 0.5 |
| 10 | 99.97 | $99.97 \pm 0.0003$ | 1.0 | 0.3667 |
| 11 | 98.23 | $98.28 \pm 0.0017$ | $< 0.01$ | 0.9969 |
| 12 | 86.15 | $86.34 \pm 0.0035$ | $< 0.01$ | 0.1666 |
| 13 | 54.27 | $54.52 \pm 0.0050$ | $< 0.01$ | $< 0.01$ |
| 14 | 22.64 | $22.78 \pm 0.0042$ | $< 0.01$ | $< 0.01$ |
| 15 | 7.29 | $7.34 \pm 0.0029$ | $< 0.01$ | 0.9235 |

| $\ell$ | Normal | Mela | $p$-value Test 1 | $p$-value Test 2 |
| --- | --- | --- | --- | --- |
| 9 | 100.0 | $99.99 \pm (< 10^{-5})$ | 1.0 | 0.5 |
| 10 | 99.97 | $99.97 \pm 0.0002$ | 1.0 | 0.1908 |
| 11 | 98.23 | $98.27 \pm 0.0015$ | $< 0.01$ | 1.0 |
| 12 | 86.15 | $86.34 \pm 0.0037$ | $< 0.01$ | 0.9581 |
| 13 | 54.27 | $54.53 \pm 0.0047$ | $< 0.01$ | $< 0.01$ |
| 14 | 22.64 | $22.78 \pm 0.0041$ | $< 0.01$ | $< 0.01$ |
| 15 | 7.29 | $7.34 \pm 0.0027$ | $< 0.01$ | 0.9399 |

#### A.2.8 Yellow fever virus indication

| $\ell$ | Normal | Lung | $p$-value Test 1 | $p$-value random |
| --- | --- | --- | --- | --- |
| 9 | 100.0 | $99.99 \pm (< 10^{-5})$ | 1.0 | 0.5 |
| 10 | 100.0 | $99.98 \pm 0.0001$ | 1.0 | 0.9805 |
| 11 | 99.05 | $99.07 \pm 0.0015$ | $< 0.01$ | 0.9999 |
| 12 | 92.10 | $92.21 \pm 0.0038$ | $< 0.01$ | 0.9061 |
| 13 | 65.28 | $65.48 \pm 0.0059$ | $< 0.01$ | 0.6362 |
| 14 | 30.54 | $30.68 \pm 0.0056$ | $< 0.01$ | 0.1009 |
| 15 | 10.81 | $10.86 \pm 0.0040$ | $< 0.01$ | 0.2397 |

| $\ell$ | Normal | Bladder | $p$-value Test 1 | $p$-value Test 2 |
| --- | --- | --- | --- | --- |
| 9 | 100.0 | $99.99 \pm (< 10^{-5})$ | 1.0 | 0.5 |
| 10 | 100.0 | $99.98 \pm 0.0001$ | 1.0 | 0.9826 |
| 11 | 99.05 | $99.07 \pm 0.0015$ | $< 0.01$ | 0.8695 |
| 12 | 92.10 | $92.21 \pm 0.0038$ | $< 0.01$ | 0.9591 |
| 13 | 65.28 | $65.47 \pm 0.0062$ | $< 0.01$ | 0.8768 |
| 14 | 30.54 | $30.67 \pm 0.0058$ | $< 0.01$ | 0.6535 |
| 15 | 10.81 | $10.85 \pm 0.0040$ | $< 0.01$ | 0.3259 |

| $\ell$ | Normal | Mela | $p$-value Test 1 | $p$-value random |
| --- | --- | --- | --- | --- |
| 9 | 100.0 | $99.99 \pm (< 10^{-5})$ | 1.0 | 0.5 |
| 10 | 100.0 | $99.98 \pm 0.0001$ | 1.0 | 0.9725 |
| 11 | 99.05 | $99.06 \pm 0.0015$ | $< 0.01$ | 0.9999 |
| 12 | 92.10 | $92.20 \pm 0.0038$ | $< 0.01$ | 0.9993 |
| 13 | 65.28 | $65.48 \pm 0.0059$ | $< 0.01$ | 0.4637 |
| 14 | 30.54 | $30.68 \pm 0.0057$ | $< 0.01$ | 0.1347 |
| 15 | 10.81 | $10.86 \pm 0.0039$ | $< 0.01$ | 0.0967 |

### A.3 Error probability data for Fig. 5

To compute 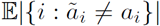 in (2) we proceeded as follows. We have 

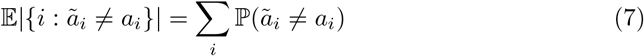

where the summation ranges over amino acid positions. Let us compute ℙ(*ã*_1_≠ *a_1_*)—for the other terms we proceed in the same way. Observe that *a_1_* is a function of the first three nucleotides *y*_1_,*y*_2_,*y*_3_ of the normal exome *y*. To emphasize this, let us write *a_1_* as *a*_1_(*y*_1_,*y*_2_,*y*_3_). Similarly, *ã*_1_ is a function of the first three nucleotides *ỹ*_1_,*ỹ*_2_,*ỹ*_3_ of the cancer genome *ỹ* and we write it as *ã*_1_(*ỹ*_1_,*ỹ*_2_,*ỹ*_3_). Therefore, we have 

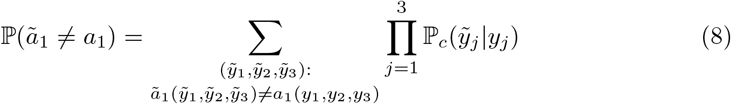

where ℙ_c_(*ỹ_j_*|*y_j_*) is the cancer channel given in the Supplementary Section A.2.

## Footnotes

1 Notice that *sℓ(x; y)* does not count multiplicity, *i.e*., strings that appear only once in the human exome and strings that appear multiple times in the human exome are counted in the same way.

2 Note that even if a cancer typically exhibits a dominant mutational signature, the simulated mutational process results in a more realistic combination of such signatures.

3 Analogously to a communication channel that alterates a transmitted message because of noise (see, *e.g*., [17]), a cancer channel alterates a DNA sequence because of somatic mutations. Hence the term “cancer channel” to emphasize the analgogy.

4 Abusing somewhat terminology, we often refer to mutation distribution as “signature.”

